# A simple, anesthesia-free infusion technique for *in vivo* metabolic tracing

**DOI:** 10.64898/2025.12.17.694756

**Authors:** Yumi Kim, Samantha Caldwell, Meredith Long, Dominique Abrahams, Margi Baldwin, Gina M. DeNicola

## Abstract

Accurate metabolic flux analysis requires tracer delivery that preserves physiological metabolism. Current methods may distort metabolism through anesthesia, surgical stress, or complex procedures. We demonstrate that isoflurane anesthesia profoundly alters serum and tissue metabolism across multiple pathways. Glycolytic and TCA cycle intermediates, sulfur and aromatic amino acid metabolites, acylcarnitines, and nucleotide pools decreased, while branched-chain amino acids, their ketoacids, ketone bodies, and fatty acids increased. These coordinated changes were suggestive of mitochondrial complex I inhibition and reduced oxidative catabolism, leading to shifts in metabolite pool sizes that compromise isotopologue-based flux interpretation. We established a tail vein catheterization method completed in minutes under brief anesthesia that enables multi-hour tracer infusion in awake, freely moving mice. This method achieved steady-state labeling of cystine and downstream products comparable to jugular infusion without supraphysiologic cystine accumulation. This platform provides a practical, physiologically accurate method for *in vivo* steady-state isotope tracing.

## Main

Metabolic regulation must coordinate energy production, macromolecule synthesis, and redox balance across multiple tissues and temporal scales to maintain organismal homeostasis^1^. This coordination becomes particularly critical during physiological challenges, including feeding, fasting, exercise or disease, when tissues must rapidly adjust their metabolic priorities while maintaining overall energy balance^1,2^. However, understanding how this inter-tissue metabolic coordination occurs requires measuring actual metabolic pathway activity rather than simply measuring metabolite levels, as the same metabolite pools can reflect vastly different flux states depending on the physiological context^3,4^.

Metabolic flux analysis addresses this limitation by quantifying the rates and directions of biochemical reactions within living systems. Stable isotope tracing has emerged as the gold standard for flux analysis *in vivo*, where labeled nutrients (e.g ^13^C, ^15^N) are introduced into an organism and their incorporation into downstream metabolites tracked by mass spectrometry or nuclear magnetic resonance^5–8^. Two distinct experimental approaches serve different experimental goals: dynamic-state tracing follows time-dependent incorporation of tracers into metabolites of interest after the start of labeling^9^, which is useful for evaluating the rates of pathways of interest; steady-state tracing maintains constant tracer levels until equilibrium, enabling estimation of the fractional nutrient contributions to downstream metabolites and pathways comparisons between tissues under physiological conditions^10^.

Several delivery routes exist for *in vivo* isotope administration. Tail vein catheterization allows for the continuous infusion of tracers under anesthesia^5^, and has been used for the study of glucose, lactate, glutamine, and other metabolites in mice^11–13^. While anesthesia may mimic metabolic studies in patients^5^, its impact on organismal and tumor metabolism may impact results. As an alternative to tracer infusion, dietary labeling is noninvasive but produces variable intake and complex relabeling patterns^14^. Jugular vein catheterization allows prolonged delivery without anesthesia, but requires complex surgery by skilled personnel and necessitates post-op recovery and analgesia^15,16^. The lack of consensus on optimal delivery methods introduces an important source of methodological variability in metabolic flux studies.

Despite the widespread use of anesthesia in *in vivo* isotope tracing studies, its effects on physiological and tissue-specific metabolism remain insufficiently characterized. Numerous studies have reported both acute and long-term metabolic effects of anesthesia. However, most have been limited to specific organs such as the brain, heart, or blood^17–20^. A physiological understanding of how anesthesia influences whole-body or organ-specific metabolism remains incomplete. In particular, the global metabolic alterations induced by isoflurane anesthesia across different tissues have not been fully investigated.

A method enabling prolonged tracer delivery without anesthesia or surgery would enhance precision, reproducibility, and accessibility of stable isotope tracing analysis *in vivo*. Here, we developed a tail vein catheter infusion system that delivers stable, prolonged isotope labeling in awake mice without the need for surgical implantation. We demonstrate that this method achieves isotopic ^13^C-cystine steady-state labeling comparable to jugular infusion while minimizing physiological stress and technical barriers, providing a practical platform for analyzing *in vivo* metabolic flux under physiological conditions.

## Results

### Effects of anesthesia on serum metabolism

To test the impact of anesthesia on metabolism, mice were subjected to anesthesia or control conditions for 2 hours, followed by serum and organ collection. Metabolites were extracted using 80% methanol containing 5mM N-ethylmaleimide (NEM), which covalently reacts with free thiols such as cysteine and glutathione, thereby preventing artifactual oxidation and allowing accurate measurement of sulfur metabolites and analyzed by LC-MS (Fig. 1A).

**Figure 1.**
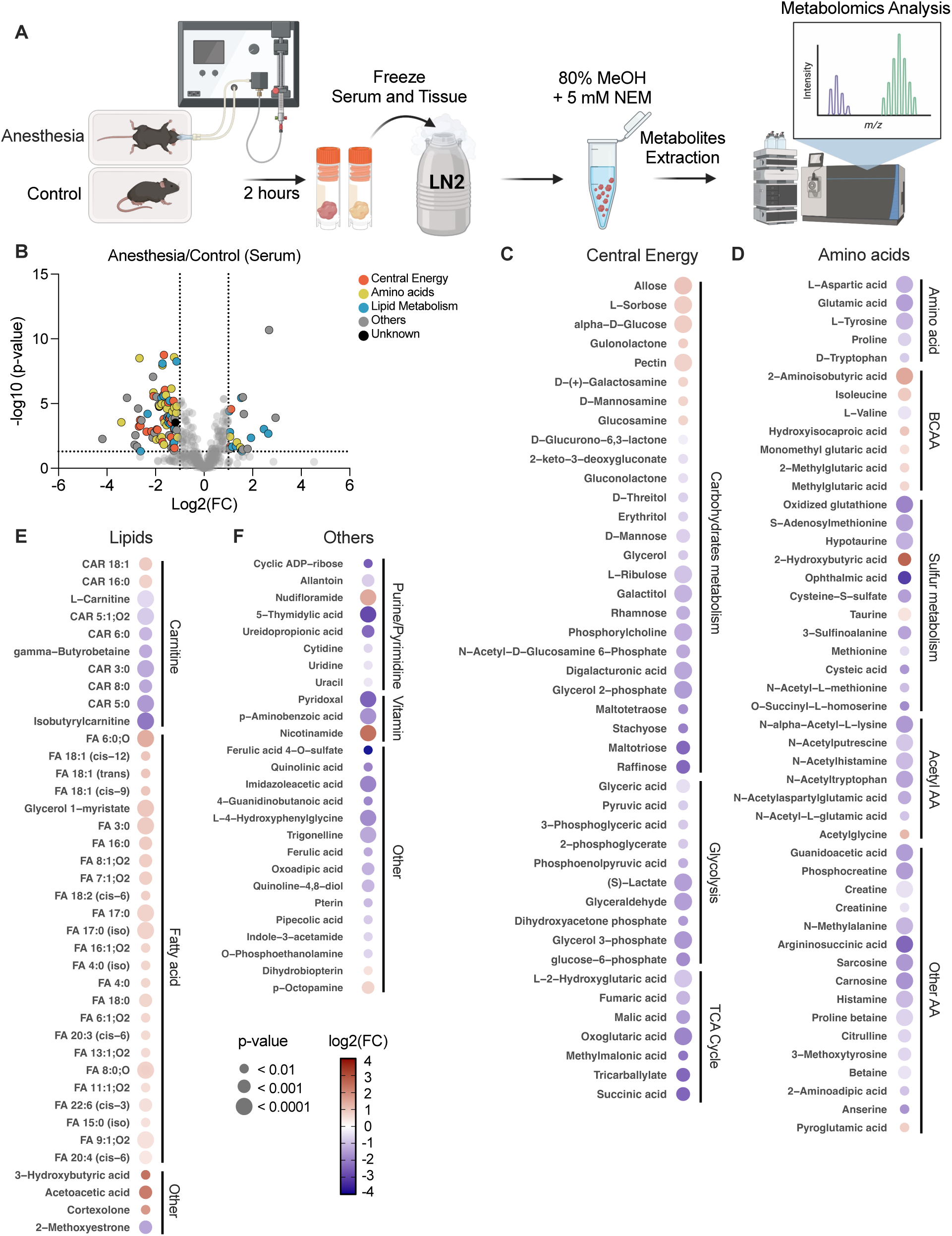
Global serum metabolomic profiling following anesthesia. (A) Experimental workflow for anesthesia and anesthesia-free conditions. Mice were subjected to either anesthesia or control conditions for 2 hours on a heating pad prior to sample collection. Serum and tissues were immediately snap-frozen in liquid nitrogen, extracted with 80% methanol containing 5 mM N-ethylmaleimide (NEM), and analyzed by LC-MS based metabolomics. (B) Volcano plot of serum metabolites colored by biochemical category (Central Energy, Amino acids/Nitrogen, Lipid Metabolism, Others, or Unknown). Dotted lines indicate significance thresholds defined as fold-change ≥ 2 and P < 0.05. (C–F) Dot-heatmaps showing significant metabolite changes within central energy, lipid, amino acid, and other metabolic pathways in anesthesia compared to anesthesia-free groups. Heatmaps display log_2_ (fold change) values for metabolites with P < 0.001 selected from volcano plot analysis. Dot size indicates P-value and dot color reflects log_2_ (fold change), with red indicating positive fold change and blue indicating negative fold change (scale −4.0 to 4.0). Data are from n = 8 mice per group.

Global metabolomics analysis revealed extensive alterations between anesthesia group and anesthesia-free control group. Among 409 serum metabolites, 102 metabolites (24.9%) showed at least a 2-fold change under anesthesia. Of these, 23 metabolites increased, and 79 metabolites decreased. Among the 23 increased metabolites, lipid metabolites represented the largest proportion (47.8%), followed by other metabolites (30.4%), amino acid metabolites (13.0%), and central energy metabolites (8.7%). Among the 79 decreased metabolites, amino acid metabolites represented the largest proportion (27.8%), followed by central energy metabolites (26.6%) and lipid metabolites (26.6%), other metabolites (16.5%), and unknown metabolites (2.5%) (Fig. 1B).

Dot heatmaps revealed coordinated metabolic alterations across central energy, amino acid, lipid, and other pathways in response to anesthesia (Fig. 1C to 1F). In the central energy category (Fig. 1C), several upstream carbohydrates, including allose, L-sorbose, and alpha-D-glucose, were increased under anesthesia, suggesting accumulation of early carbohydrate intermediates. In contrast, multiple downstream carbohydrate metabolites displayed strong reductions with high statistical significance. Raffinose, maltotriose, digalacturonic acid, glycerol-2-phosphate, phosphorylcholine, and N-acetyl-D-glucosamine-6-phosphate exhibited the largest decreases. Downstream glycolytic and tricarboxylic acid intermediates were consistently decreased by anesthesia. Metabolites such as 3-phosphoglyceric acid, glyceraldehyde, lactate, succinic acid, fumaric acid, malic acid, and oxoglutaric acid were all decreased upon anesthesia, indicating coordinated suppression of glycolytic throughput and TCA cycle activity, resulting in the accumulation of upstream carbohydrates.

Amino acid metabolism showed broad suppression under anesthesia (Fig. 1D). The strongest reductions were observed in oxidized glutathione and ophthalmic acid, both products of the glutathione synthesis machinery. Additional sulfur-related metabolites, including S-adenosylmethionine, cysteine-S-sulfate, sulfinoalanine, hypotaurine, and cysteic acid, were also decreased, reflecting coordinated reductions across sulfur-linked amino acid pathways. In contrast, 2-hydroxybutyric acid showed one of the largest increases within this category. This metabolite is a well-characterized product of 2-ketobutyrate reduction and arises from established pathways involving threonine and methionine metabolism. Its strong elevation indicates a selective increase in upstream sulfur-associated intermediates despite reductions in downstream sulfur utilization. Most proteinogenic amino acids exhibited similar reductions. Aspartate, glutamic acid, tyrosine, tryptophan, histidine, and several acetylated amino acids, including N-acetyl-L-glutamic acid and N-acetyl-L-lysine, were significantly decreased. Despite these overall decreases, several metabolites increased, including isoleucine, hydroxyisocaproic acid, and 2-aminoisobutyric acid. These metabolites belong to well-established branched-chain and modified amino acid pathways, suggesting selective preservation or accumulation of these intermediates under anesthesia. Together, these findings show that anesthesia broadly suppresses amino acid and sulfur metabolism while selectively increasing 2-hydroxybutyric acid and several branched-chain–associated metabolites.

Lipid metabolism showed distinct and pathway-specific changes under anesthesia (Fig. 1E). The most noted decreases were observed in acylcarnitine species, including CAR 5:0, CAR 6:0, CAR 8:0, CAR 3:0, and isobutyrylcarnitine, which displayed the largest negative fold changes with high statistical significance. These metabolites are well-established intermediates involved in fatty-acid transport into mitochondria, and their strong reductions indicate a marked suppression of carnitine-linked lipid handling during anesthesia. In contrast, numerous fatty acids exhibited the strongest increases within the lipid category. These increases spanned short-, medium-, and long-chain species, ranging from FA 4:0 to FA 22:6. Both saturated fatty acids such as FA 16:0 and FA 17:0 and unsaturated fatty acids including FA 18:1 (cis-9), FA 18:1 (cis-12), FA 20:3, and FA 22:6 showed significant elevations. This pattern indicates a broad rise in circulating fatty-acid pools of multiple chain lengths and saturation states under anesthesia. Ketone-associated metabolites showed consistent elevations under anesthesia. Both 3-hydroxybutyric acid and acetoacetate were strongly increased, reflecting increased production of well-established ketone intermediates. In contrast to these increases, 2-methoxyestrone showed a significant decrease, representing one of the few lipid-associated metabolites reduced outside the acylcarnitine group. Additional metabolites, including several nucleotide- and vitamin-associated compounds, also showed changes under anesthesia, although these patterns were less coordinated than those observed in the major metabolic groups (Fig. 1F). Together, these findings show that anesthesia suppresses acylcarnitine-dependent fatty-acid transport while selectively increasing multiple fatty acids and ketone-related metabolites.

### Organ-specific metabolic alterations induced by anesthesia

The changes in serum metabolites may result from disrupted metabolism in metabolically active organs, or they may alter the delivery of substrates to these organs, which in turn affects their metabolic processes. To this end, we conducted metabolite profiling across four organs, which revealed both shared and tissue-specific metabolic alterations induced by anesthesia (Fig. 2A to 2D). In the brain (Fig. 2A), the strongest reductions centered on carbohydrate-linked and carnitine-linked intermediates, including galactitol and isobutyrylcarnitine, indicating attenuation of polyol metabolism and short-chain acyl-carnitine turnover. In contrast, increases in leucine and its transamination product, alpha-ketoisocaproic acid, suggest an elevation in branched-chain amino acid (BCAA) catabolism.

**Figure 2.**
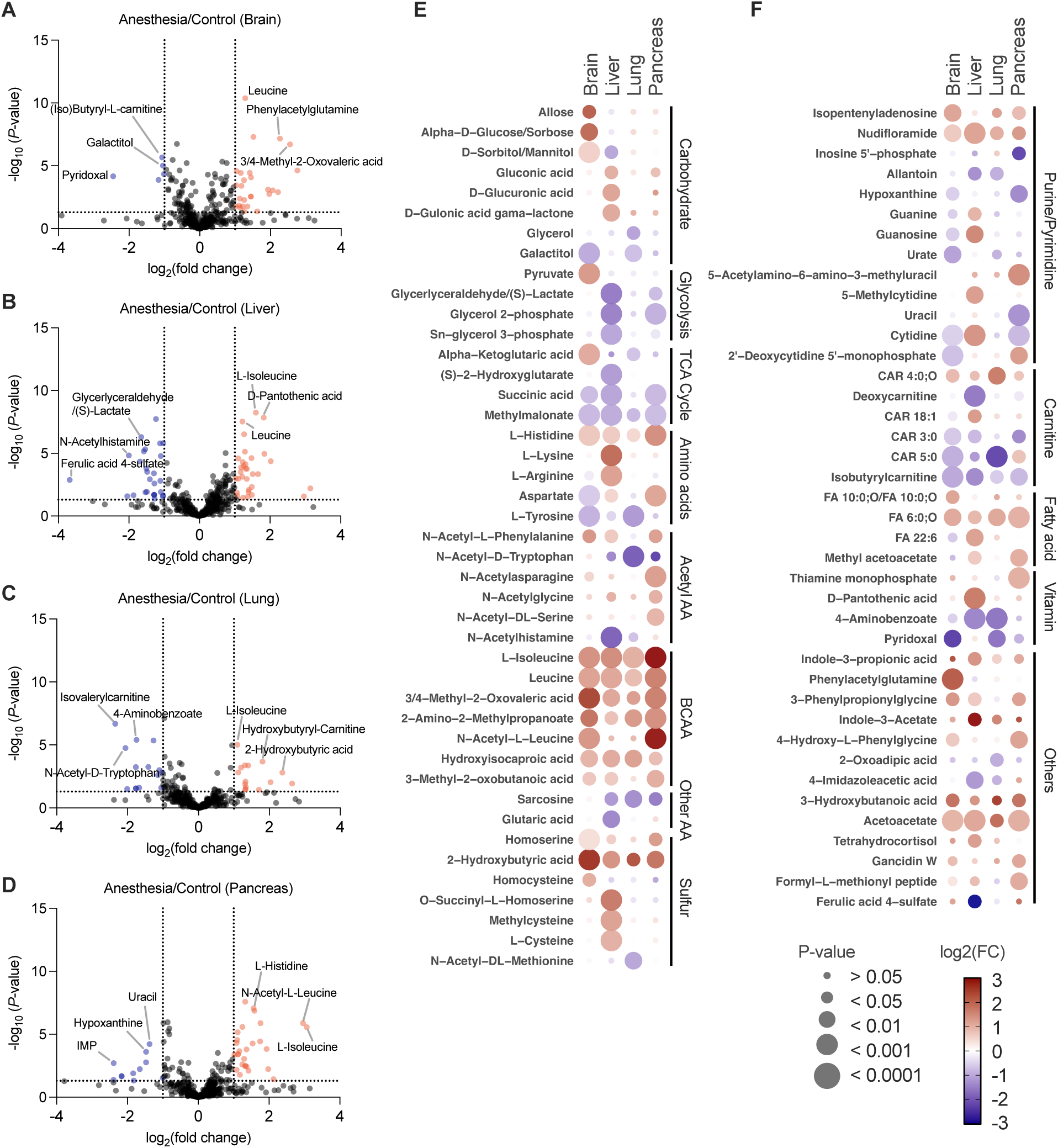
Metabolomic changes across brain, lung, liver, and pancreas induced by anesthesia. Metabolite alterations in brain, lung, liver, and pancreas after 2 hours of anesthesia versus control. Metabolites were profiled by LC-MS. (A–D) Volcano plots of global metabolomics in brain (A), lung (B), liver (C), and pancreas (D). Metabolites are shown in red (up-regulated) and blue (down-regulated), with selected representative metabolites labeled. Plots display log_2_ (fold change) versus −log_10_ (P-value) for anesthesia compared to anesthesia-free groups. (E and F) Dot-heatmap showing metabolic changes grouped by biochemical pathway across all four tissues. For each tissue, metabolites that passed the cutoff in the corresponding volcano plot (fold-change ≥ 2 and P < 0.01, unpaired Student’s t-test) were selected. Dot size indicates P-value and dot color represents log_2_ (fold change) in anesthesia compared to control (scale −3 to 3). Data are from n = 8 mice per group.

The liver displayed the broadest metabolic remodeling across categories (Fig. 2B). Strong decreases in glyceraldehyde and lactate suggested reduced glycolytic flux, while reductions in N-acetylhistamine and ferulic acid 4-sulfate indicated suppression of nitrogen-associated and phenolic pathways. At the same time, increases in leucine, isoleucine, and pantothenic acid highlighted shifts toward amino-acid–derived carbon flux and CoA-associated metabolism.

In the lung (Fig. 2C), marked decreases in 4-aminobenzoate, isovalerylcarnitine, and N-acetyl-D-tryptophan indicated reductions in vitamin-related pathways, short-chain acyl-carnitines, and aromatic amino acid derivatives. Conversely, increases in isoleucine, 2-hydroxybutyric acid, and hydroxybutyrylcarnitine suggested enhancement of BCAA–related and hydroxy-acid pathways.

The pancreas showed the largest decreases in nucleobase-associated metabolites, including uracil, hypoxanthine, and inosine 5′-phosphate (Fig. 2D), indicating suppression of purine and pyrimidine metabolism. In contrast, histidine, isoleucine, and N-acetyl-leucine were among the most strongly increased metabolites, pointing toward selective activation of cationic amino acid and BCAA metabolism.

Pathway-level analysis from the dot-heatmap revealed biochemical signatures that were consistent across all examined tissues (Fig. 2E and 2F). BCAA and their corresponding alpha-ketoacids, including leucine, isoleucine, 3-methyl-2-oxobutanoic acid, and 3 or 4-methyl-2-oxovaleric acid, were uniformly increased, indicating a systemic shift toward enhanced BCAA catabolism (Fig. 2E). Ketone-associated metabolites, such as 3-hydroxybutyrate and acetoacetate, also increased across multiple organs. Short- and medium-chain acyl-carnitines, including isobutyrylcarnitine, CAR 3:0, and CAR 5:0, consistently decreased, reflecting suppression of fatty acid transport through the carnitine shuttle (Fig. 2F). Succinic acid and methylmalonate, both TCA cycle intermediates, were decreased in brain, liver, lung, and pancreas. In amino acid metabolism, histidine increased, and tyrosine decreased across the same tissues. In sulfur metabolism, 2-hydroxybutyric acid increased in all organs, whereas other sulfur metabolites changed primarily in the liver. Most pathway-level alterations, including glycolysis, the TCA cycle, amino acids, and sulfur metabolites, were primarily observed in the liver. Acetylated amino acids changed primarily in the pancreas, with N-acetylphenylalanine, N-acetylasparagine, N-acetylglycine, and N-acetylserine increasing only in this organ. In contrast, N-acetyltryptophan decreased in the pancreas, liver, and lung.

Overall, these findings demonstrate that anesthesia induces a coordinated metabolic remodeling across serum and multiple organs. Several serum alterations parallel the tissue-level changes, including decreases in acyl-carnitines and amino acids and increases in BCAA-related metabolites. These correspondences suggest that reduced hepatic fatty acid oxidation and altered amino acid metabolism in the liver and other tissues may contribute to the circulating metabolic patterns observed under anesthesia. At the same time, some serum changes may reflect multi-organ integration of metabolic adjustments rather than originating from a single tissue. Together, these data indicate that anesthesia produces a systemic shift in metabolic homeostasis that is reflected across both serum and organ compartments.

### Tail vein infusion provides physiological tracer delivery

Stable isotope infusion in mice has traditionally been performed either through tail vein infusions under anesthesia or through surgically implanted jugular catheters. Both approaches introduce metabolic disturbances, either through anesthetic exposure or through surgical stress and postoperative recovery, and they also present technical challenges that limit reproducibility and their broad applicability across research labs. To overcome these limitations, we developed a tail vein catheter infusion method that enables stable isotope delivery on conscious mice without the need for surgical implantation.

We compared jugular vein and tail vein catheterization as delivery routes for U-^13^C_6_-cystine infusion, which we previously established for *in vivo* isotope tracing^21^. Jugular catheterization required a surgical procedure with 30 to 40 minutes of operative time, prolonged anesthesia, 2 to 5 days of postoperative recovery, and mandatory pain medication (Fig. 3A). In contrast, tail vein catheterization was a non-surgical approach completed within 5 to 10 minutes under brief anesthesia and required no recovery period or pain management (Fig. 3B and 3C). We established a standardized tail vein catheterization workflow for anesthesia-free infusion studies. The procedure began by fitting mice with a harness through which the external infusion line from the pump was threaded. A 27 G catheter needle was then inserted into the lateral tail vein (Fig. 3B), secured with a small drop of cyanoacrylate adhesive, and reinforced with surgical tape to protect and stabilize the intravenous access point (Fig. 3C). A temporary subcutaneous tunnel was created by inserting a 16 G introducer catheter through the dorsal skin and advancing it subcutaneously toward the base of the tail. This tunnel allowed the external infusion line to be guided from the dorsal entry point through the subcutaneous tract and out near the tail base. Once the infusion line emerged at the tail base, the introducer was removed and the infusion line was connected to the tail vein catheter via a blunted connector, as shown in Fig. 3C. After connection, the line was gently pulled by hand so that the junction was drawn into the harness and housed within a protective metal spring tether before being attached to the programmable syringe pump. Once the catheter infusion setup is completed, mice are returned to their cages and connected to a pump system that delivered the isotope tracer at a constant rate for the designated infusion period (Fig. 3D). Mice recovered rapidly and moved freely during infusion, as shown in Supplementary Video 1, indicating minimal physiological disturbance.

**Figure 3.**
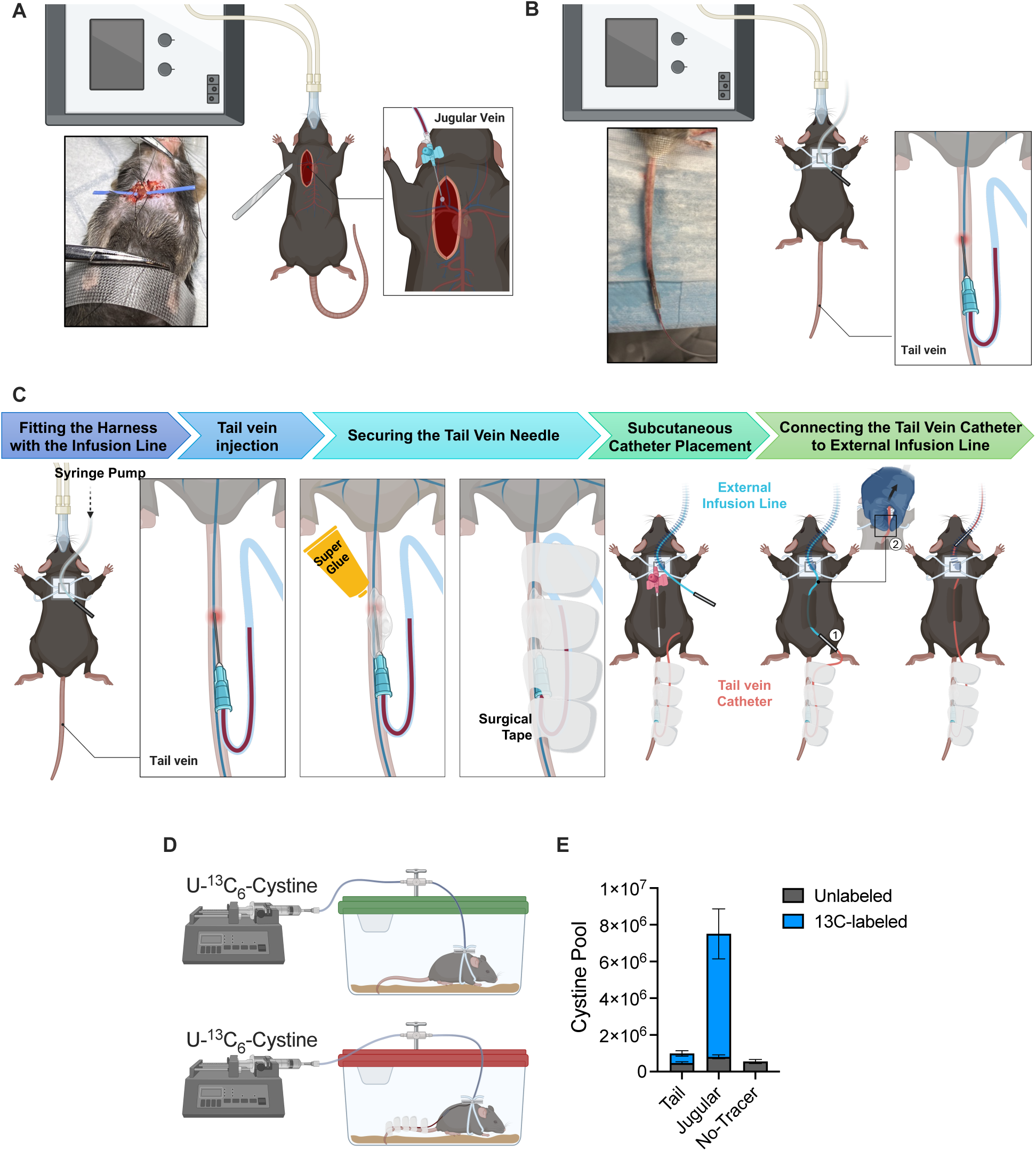
Comparison of jugular and tail vein catheterization for U-^13^C_6_-cystine infusion. (A–B) Schematic diagrams of (A) jugular vein catheterization, which requires a surgical procedure under prolonged anesthesia, and (B) tail vein catheterization, which is performed under brief anesthesia without surgery. (C) Workflow of the tail vein catheter infusion setup. The external infusion line connected to the syringe pump is threaded through the mouse harness, and the harness is fitted onto the mouse. The tail vein catheter is inserted into the lateral tail vein, and the needle is stabilized using cyanoacrylate adhesive followed by surgical tape reinforcement. After needle fixation, a subcutaneous tunnel is created by advancing a catheter introducer through the dorsal skin toward the base of the tail. (1) The external infusion line is then guided through this tunnel and connected to the tail vein catheter via a blunted connector. (2) Finally, the line is gently pulled to draw the junction into position, completing the infusion circuit. (D) Experimental setup for tracer delivery. After catheter placement and recovery, 10 mg/mL U-^13^C_6_-cystine was infused continuously using an automated pump at a rate of 120 μL per hour for 4 hours. At the end of infusion, serum and tissues were collected, snap-frozen in liquid nitrogen, and processed for metabolomics. Serum was extracted with 80% methanol containing 5 mM N-ethylmaleimide to stabilize sulfur metabolites and analyzed by LC-MS in positive ion mode. (E) Total cystine pool size in serum, showing unlabeled and ^13^C-labeled cystine in jugular, tail, and no-tracer groups. Data are mean ± SEM (n = 12 for jugular, n = 13 for tail, n = 8 for no-tracer).

Measurement of the cystine pool showed that jugular infusion markedly elevated the total cystine pool compared with tail infusion or no-tracer controls. In contrast, tail vein infusion maintained circulating cystine levels close to the physiological baseline (Fig. 3E).

Together, these findings indicate that while both approaches support reliable isotopologue analysis, tail vein catheterization provides a minimally invasive, low-risk, and physiologically faithful method for stable isotope infusion. Compared with jugular catheterization, the tail approach reduces procedural burden, eliminates surgical recovery, and maintains endogenous metabolite pool sizes while still enabling reliable flux measurements.

### Tail vein infusion provides stable labeling of cystine-derived metabolites in tissues

The schematic illustrates the incorporation of U-^13^C_6_-cystine into downstream metabolites, including cysteine, γ-glutamylcysteine, glutathione, hypotaurine, and taurine (Fig. 4A). Red carbon atoms indicate the expected ^13^C labeling patterns, with U-^13^C_6_-cystine generating distinct isotopologue species (M+2, M+3, and M+6) that enable quantitative tracing of cystine utilization.

**Figure 4.**
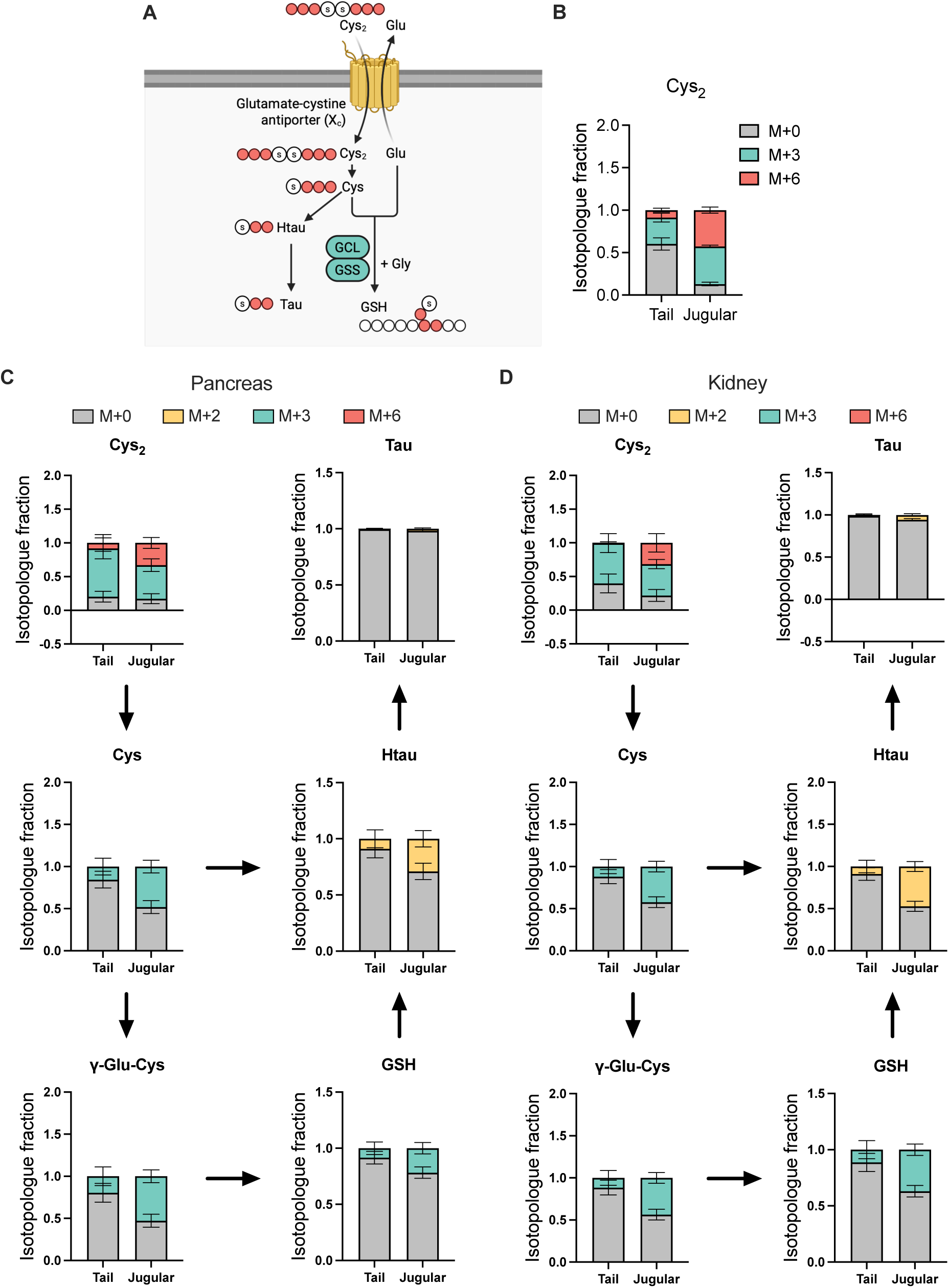
Isotopologue tracing of cystine incorporation into glutathione and taurine pathways. (A) Schematic of cystine utilization pathway, illustrating incorporation into cysteine, γ-glutamylcysteine, glutathione, hypotaurine, and taurine. Labeled carbon atoms from U-^13^C_6_-cystine are indicated in red. (B) (B–D) Isotopologue analysis of serum, pancreas, and kidney collected after U-^13^C_6_-cystine infusion via jugular or tail vein catheterization. (B) Serum cystine isotopologue distribution (M+0, M+3, M+6). (C–D) Isotopologue fractions of cystine-derived metabolites in pancreas (C) and kidney (D), showing incorporation into cysteine, γ-glutamylcysteine, glutathione, hypotaurine, and taurine. Data are presented as mean ± SEM (n = 12 for jugular, n = 13 for tail).

To evaluate tracer labeling achieved by tail vein versus jugular vein catheterization, serum and tissues were collected immediately after the infusion period, snap frozen in liquid nitrogen, stored at −80°C, and later extracted with 80% methanol containing 5 mM NEM prior to LC-MS analysis. In serum, isotopologue analysis of circulating cystine showed that tail and jugular infusion produced different degrees of labeling but similar isotopologue hierarchies (Fig. 4B). Tail infusion maintained a substantial unlabeled fraction, with total ^13^C labeling (M+3 and M+6) of roughly 44%, whereas jugular infusion drove cystine labeling to nearly 87%. Thus, while jugular infusion drives higher tracer pool expansion, tail vein catheterization achieves steady-state labeling with minimal perturbation of endogenous pools.

In pancreas and kidney, tissue cystine (Cys_2_) was highly labeled in both conditions, with total labeled fractions close to 80% for tail and jugular infusion alike (Fig. 4C and 4D). This indicated that the infused tracer efficiently equilibrated with tissue cystine pools regardless of infusion route. For downstream metabolites, including cysteine, γ-glutamylcysteine, glutathione, hypotaurine, and taurine, the overall isotopologue patterns were similar between routes, although fractional labeling was modestly lower after tail infusion than after jugular infusion. Despite this quantitative difference, all metabolites showed clear incorporation of ^13^C from U-^13^C_6_-cystine in both tissues, confirming that the tracer robustly reports flux through glutathione and taurine biosynthetic pathways.

Taken together with the pool-size data, these results show that tail vein infusion achieves strong and informative labeling of cystine and its downstream metabolites while maintaining circulating cystine pools closer to physiological levels. Jugular infusion yields higher fractional labeling of circulating cystine but at the cost of supraphysiologic pool expansion and the need for surgical catheterization, whereas tail vein infusion provides a technically simpler and more physiologically faithful alternative that still supports reliable isotopologue-based flux analysis.

## Discussion

As *in vivo* metabolomics and stable isotope tracing continue to expand, achieving physiologically accurate measurements in mouse models has become increasingly important^8,22^. These experimental approaches have traditionally relied on dietary labeling^14,23^, tail vein infusions performed under continuous anesthesia^17,19,20^, or surgical jugular catheterization for prolonged tracer delivery^24–26^. However, each of these approaches introduces physiological or technical variables that complicate accurate interpretation of *in vivo* flux. Dietary labeling results in uncontrolled nutrient intake and complex relabeling patterns. Tail vein delivery has also been used for steady-state isotope infusion, but these procedures are performed under continuous anesthesia^5^, which introduces physiological disturbances and alters systemic metabolism. Jugular catheter infusion can achieve steady state labeling but necessitates surgery, prolonged anesthesia, external thermoregulation to prevent hypothermia, postoperative recovery, and analgesic administration, all of which introduce physiological perturbations^15,16^. Despite the widespread use of anesthesia in these procedures, its organ level metabolic effects remain incompletely characterized. Here, we systematically define how isoflurane anesthesia reprograms systemic metabolism and establish a nonsurgical tail vein catheter infusion platform that enables physiological steady state isotope tracing on conscious animals, while minimizing procedural and metabolic disruption.

To investigate whether anesthesia alters metabolism, we analyzed serum and multiple tissues collected from mice exposed to anesthesia or control anesthesia-free conditions. Isoflurane broadly inhibited central carbon metabolism and amino acid pathways. Glycolytic and tricarboxylic acid related metabolites were reduced, as were methionine based one carbon intermediates and sulfur metabolites, accompanied by a general decrease in amino acids. In contrast, BCAA and their corresponding alpha ketoacids accumulated, indicating a shift in amino acid handling. Reductions in short and medium chain acylcarnitines together with elevations in ketone related metabolites suggested inhibition of fatty acid oxidation and diversion toward alternative fuel usage. Similar signatures were reproduced across multiple organs. The brain exhibited increases in sugars and energy related intermediates, the lung demonstrated elevated glycolytic intermediates and reductions in aromatic amino acids, the liver showed increases in indole related metabolites and decreases in purine and pyrimidine synthesis intermediates, and the pancreas displayed elevated BCAAs and ketones with reduced nucleotide intermediates. These coordinated, yet tissue-specific, signatures indicate that anesthesia induces a distinct metabolic state that extends beyond reduced energy demand and alters multiple carbon utilization and amino acid processing pathways simultaneously.

The metabolic rewiring induced by anesthesia aligns with known mechanisms of isoflurane action and with previous reports documenting anesthesia associated metabolic perturbations. Isoflurane inhibits mitochondrial complex I, decreasing oxygen consumption and ATP production, consistent with reductions in oxidative metabolism and fatty acid oxidation observed across tissues^27,28^. Suppressed BCAA oxidation and consequent BCAA accumulation have been reported in muscle and liver and match the elevations detected in our study^29^. Acute alterations in serum glucose, lactate, and non-esterified fatty acids observed here are consistent with findings showing that diverse anesthetics and sampling procedures immediately distort circulating metabolites^30^. Isoflurane induced increases in brain lactate and glutamate and metabolic patterns distinct from propofol anesthesia^17^ parallel the brain specific shifts identified in our dataset and underscore that anesthesia produces anesthetic specific metabolic programs. Additional studies comparing awake versus isoflurane anesthesia conditions in the prefrontal cortex similarly reported decreased amino acids and increased nucleosides under anesthesia^19^, reinforcing that anesthesia represents a metabolic state distinct from wakefulness. Furthermore, isoflurane has been reported to induce not only acute but also long-lasting transcriptomic remodeling in the heart, particularly maladaptive metabolic reprogramming in aged animals^18^, supporting the notion that anesthesia is an active metabolic stressor rather than a neutral baseline condition. Collectively, these findings demonstrate that anesthesia alters pool sizes, substrate availability, amino acid and fatty acid utilization, and redox state, all of which directly influence isotopologue distributions and can distort flux interpretation. Minimizing anesthetic exposure is therefore essential for accurately assessing physiological *in vivo* flux.

To address this need, we developed a nonsurgical tail vein catheter infusion system that supports prolonged tracer delivery under physiological conditions in conscious animals. Catheter placement required only five to ten minutes of anesthesia, after which mice recovered immediately and moved freely throughout infusion. Unlike conventional tail vein catheterization and surgical jugular catheterization, this approach avoids extended anesthesia, analgesic administration, and recovery time, thereby reducing physiological disturbance and improving accessibility. Maintaining natural behavioral and metabolic states during infusion lowers environmental constraints on flux measurements and supports the acquisition of physiologically relevant metabolic data.

Using U-^13^C_6_-cystine infusion, we evaluated the performance of this platform and found that fractional isotopologue labeling was comparable to jugular catheter infusion, yet the tracer pool size differed markedly. Jugular infusion increased the circulating cystine pool above physiological levels, whereas awake tail infusion-maintained pool sizes similar to those in non-tracer controls. This distinction is critical because supraphysiological tracer spikes can distort endogenous flux relationships The difference likely reflects delivery kinetics, with jugular infusion introducing tracer rapidly into central circulation, while tail infusion delivers tracer more gradually, potentially leading to increased tracer clearance and lower tracer pool size. Despite decreased tracer delivery, downstream cysteine, γ-glutamylcysteine, glutathione, hypotaurine, and taurine exhibited stable and consistent labeling patterns during tail vein infusion, supporting reliable pathway level flux estimation. Thus, tail vein infusion on conscious mice preserves physiological metabolic conditions while enabling steady state labeling, addressing key limitations of existing surgical approaches and improving the accuracy of steady state flux analysis.

In summary, this study provides a multi organ characterization of how isoflurane anesthesia reprograms systemic metabolism and demonstrates that nonsurgical tail vein catheter infusion on conscious mice enables physiological steady state isotope tracing *in vivo*. By avoiding anesthesia induced metabolic distortion while maintaining robust and reproducible labeling, this platform offers a practical and metabolically accurate method for *in vivo* flux analysis and is broadly applicable to diverse physiological and disease models.

## Methods

### Animal care

All animal experiments were conducted in accordance with the ethical guidelines and approval of the University of South Florida Institutional Animal Care and Use Committee (Protocols IS00012540R and IS00008736R). C57BL/6NJ mice (Jackson Laboratory, Strain #005304) were housed in the Animal Research Facility at the Moffitt Cancer Center under specific pathogen-free conditions with a 12-h light/dark cycle (lights on at 6 AM and off at 6 PM), ambient temperature of 21–23 °C, and relative humidity maintained at 30–70%. Mice had ad libitum access to standard chow (Envigo Bioproducts Inc T.2918.CS) and water.

### Anesthesia

C57BL/6NJ mice were used for anesthesia and anesthesia-free comparison experiments. Mice were randomly assigned to either an anesthesia group or a control group. For the anesthesia condition, mice were exposed to 2% isoflurane delivered via an induction chamber for 2 hours. The anesthesia-free group were littermate controls collected without exposure to isoflurane. All mice were continuously monitored to ensure normal respiration and to minimize stress. At the end of the 2-hour period, blood was first collected from the submandibular vein, followed by euthanasia by cervical dislocation. Serum and tissues were rapidly collected, placed in pre-labeled cryovials, and flash-frozen in liquid nitrogen for subsequent LC–MS analysis.

### Stable isotope animal infusions

To prepare the ^13^C_6_-cystine tracer, ^13^C_3_-cysteine (10 mg/mL in sterile saline; Cambridge Isotope Laboratories, CLM-4320-H-PK) was oxidized with an equimolar quantity of H_2_O_2_ for 30 minutes at room temperature, and the reaction was terminated by heating at 60 °C for 5 minutes. The resulting cystine was resolubilized with 6 N HCl before infusion. Jugular vein catheterization and stable isotope infusions were performed as previously described^21^. Briefly, indwelling catheters were surgically implanted into the right jugular vein 2–7 days before infusion, and mice were allowed to recover under standard housing conditions. On the day of infusion, catheters were connected to a syringe pump through a swivel and tether system (SAI Infusion Technologies) that allowed free movement within the cage. A spring-supported harness was used to secure the tether and prevent disconnection.

For tail vein catheter infusions, the infusion line was assembled using a 3 mL syringe (Monoject™ SoftPack Syringe, Covidien 1180300777) fitted with a 25 G needle (Exchange Supplies, Code:TO25) connected to pre-cut polyethylene tubing, and joined with blunted 25 G connectors to ensure continuous flow. The line was filled with tracer solution and checked for patency prior to use. The completed line was routed through a stainless-steel spring tether. Mice were fitted with a harness, anesthetized with 2% isoflurane, and placed on a heating pad to facilitate vasodilation. A 27 G tail vein catheter with butterfly tabs (SAI Infusion Technologies, BF-27-01) removed was inserted into the lateral tail vein and flushed with 0.3–0.4 mL of sterile saline to confirm placement. The catheter was fixed with tissue adhesive and secured with surgical tape along the length of the tail. To externalize the infusion line, a 16 G Surflash polyurethane I.V. catheter (Terumo, SC-360106,) was used to tunnel the tubing subcutaneously from the base of the tail to the mid-back, exiting beneath the harness connection point. The tail catheter was then attached to the infusion line via a blunted 25 G connector, drawn into a protective metal spring tether, and connected to a programmable syringe pump (BS-8000, Braintree Scientific). As with jugular catheter infusions, the tubing and pump were integrated into a swivel and spring-supported tether system (SAI Infusion Technologies) that allowed mice to move freely within the cage throughout the infusion period.

All mice that had been set up for infusion received 10 mg/mL ^13^C_6_-cystine at a constant rate of 120 μL/hour for 4 hours. During the final minutes of infusion, blood was collected from the submandibular vein, and mice were euthanized by cervical dislocation. Tissues of interest were rapidly excised, placed in cryovials, and flash-frozen in liquid nitrogen for subsequent LC–MS analysis.

### Sample preparations

Serum and tissue samples were extracted using 80% methanol-based solvent containing 5 mM N-ethylmaleimide (NEM) prepared in 10 mM ammonium formate (pH 7.0). For anesthesia and anesthesia-free samples, internal standards (Cambridge Isotope Laboratories, Inc., Cat# MSK-CAA) were included in the extraction solvent. For stable isotope infusion samples, the extraction solvent was identical except that internal standards were omitted. Frozen serum aliquots (10 μL) were thawed on ice, mixed with 90 μL of ice-cold extraction solvent by pipetting to fully resuspend the sample. The mixtures were then stored at −80 °C overnight. Frozen tissues were pulverized under liquid nitrogen using a pre-chilled BioPulverizer (BioSpec). Pulverized tissue (50 mg per mL extraction solvent) was transferred into pre-cooled tubes, mixed with 80% methanol containing 5 mM NEM in 10 mM ammonium formate (pH 7.0), and stored at −80 °C overnight. The following day, all samples were centrifuged at 17,000 × g for 20 minutes at 4 °C, and clarified supernatants were transferred to LC–MS vials and stored at −80 °C until analysis.

### LC-MS analysis and data processing

Metabolite analysis was performed using a Vanquish UPLC system (Thermo Fisher Scientific, Waltham, MA) coupled to a Q-Exactive HF Orbitrap mass spectrometer equipped with a heated electrospray ionization (HESI) source (Thermo Fisher Scientific). Separation was achieved on an Atlantis Premier BEH Z-HILIC VanGuard FIT column (2.1 × 150 mm, 2.5 µm; Waters, Milford, MA) maintained at 30 °C. The mobile phases consisted of solvent A (10 mM ammonium carbonate and 0.05% ammonium hydroxide in water) and solvent B (100% acetonitrile). The chromatographic gradient was programmed as follows: 0 min, 80% B; 13 min, 20% B; 15 min, 20% B; followed by re-equilibration to initial conditions. The flow rate was 150 µL/min, and the injection volume was 5 µL.

The Q-Exactive HF was operated in positive and negative electrospray ionization (ESI) modes, acquired in separate runs. The MS1 scan range was m/z 65–975 for negative mode and m/z 65–950 for positive mode. Full MS scans were collected at a resolution of 120,000 (m/z 200) with an automatic gain control (AGC) target of 3 × 10^6^.

Raw data files (.raw) were converted to .cdf format using Xcalibur (Version 4.0), and further processed using EI-MAVEN (Version 0.12.0). Data processing was performed with default parameters except as follows: ionization mode (positive or negative), isotopic tracer (13C), and extracted ion chromatogram (EIC) extraction window ±10 ppm. Metabolite identification was based on accurate mass and retention time matched to an established in-house library of authentic standards (Tsugawa et al., Nature Methods, 2015). Peak intensities were measured as AreaTop (mean of the three highest points of each EIC). Data from isotope labeling experiments was corrected for natural abundance using IsoCor 13C. Absolute quantitation of serum and tissues metabolites was performed using internal standard labeled with stable isotopes.

## Author Contribution

Y.K. and G.M.D. conceived the study and designed experiments. Y.K. performed metabolomics analyses and interpreted the data. S.C. and M.L. performed catheter infusions and assisted with animal care and tissue collection. M.B. and D.A. helped establish and optimize the tail-vein catheter infusion method. G.M.D. supervised the project and acquired funding. Y.K. wrote the manuscript, and all authors discussed the results and commented on the manuscript.

## Acknowledgements

We would like to thank Aimee Falzone for assisting with catheter infusions for this study and the members of the DeNicola laboratory for their valuable discussions. This work was supported by the NIH/NCI (P01CA250984) to G.M.D. We also acknowledge the support of the Comparative Medicine and Proteomics/Metabolomics Core Facilities at the H. Lee Moffitt Cancer Center & Research Institute which are funded in part by Moffitt’s Cancer Center Support Grant (NCI, P30-CA076292). All schematics were created with BioRender.com.

## Supplementary Information

### Supplementary Table

**Supplementary Table 1. Serum metabolite changes under anesthesia.** Normalized metabolomics data comparing serum samples from anesthesia and anesthesia-free conditions, including fold changes and p-values.

**Supplementary Table 2. Brain metabolite changes under anesthesia.** Normalized metabolomics data from brain tissue comparing anesthesia and anesthesia-free conditions, including fold changes and p-values.

**Supplementary Table 3. Liver metabolite changes under anesthesia.** Normalized metabolomics analysis of liver tissue comparing anesthesia and anesthesia-free conditions, including fold changes and p-values.

**Supplementary Table 4. Lung metabolite changes under anesthesia.** Metabolite-level changes in lung tissue under anesthesia relative to anesthesia-free conditions, based on normalized metabolomics data with associated fold changes and p-values.

**Supplementary Table 5. Pancreatic metabolite changes under anesthesia.** Metabolite changes in pancreas tissue under anesthesia compared with anesthesia-free conditions, derived from normalized metabolomics data.

### Supplementary Video

**Supplementary Video 1. Tail-vein catheter infusion in awake, freely moving mice** This video shows a mouse after tail-vein catheter placement and transfer to the infusion cage. During continuous isotope tracer infusion, the mouse moves freely without restraint, demonstrating stable behavior and minimal physiological disturbance throughout the procedure.

